# The Precision and Power of Population Branch Statistics in Identifying the Genomic Signatures of Local Adaptation

**DOI:** 10.1101/2024.05.14.594139

**Authors:** Max Shpak, Kadee N. Lawrence, John E. Pool

## Abstract

Population branch statistics, which estimate the branch lengths of focal populations with respect to two outgroups, have been used as an alternative to F_ST_-based genome-wide scans for identifying loci associated with local selective sweeps. In addition to the original population branch statistic (PBS), there are subsequently proposed branch rescalings: normalized population branch statistic (PBSn1), which adjusts focal branch length with respect to outgroup branch lengths at the same locus, and population branch excess (PBE), which also incorporates median branch lengths at other loci. PBSn1 and PBE have been proposed to be less sensitive to allele frequency divergence generated by background selection or geographically ubiquitous positive selection rather than local selective sweeps. However, the accuracy and statistical power of branch statistics have not been systematically assessed. To do so, we simulate genomes in representative large and small populations with varying proportions of sites evolving under genetic drift or background selection (approximated using variable *N*_e_), local selective sweeps, and geographically parallel selective sweeps. We then assess the probability that local selective sweep loci are correctly identified as outliers by F_ST_ and by each of the branch statistics. We find that branch statistics consistently outperform F_ST_ at identifying local sweeps. When background selection and/or parallel sweeps are introduced, PBSn1 and especially PBE correctly identify local sweeps among their top outliers at a higher frequency than PBS. These results validate the greater specificity of rescaled branch statistics such as PBE to detect population-specific positive selection, supporting their use in genomic studies focused on local adaptation.

**Significance Statement:** Population branch statistics are widely used in genome-wide scans to identify loci associated with local adaptation. This study finds that branch statistics are more accurate than *F*_ST_ at identifying local selective sweeps under a wide range of demographic parameters and models of evolution. It also demonstrates that certain branch statistics have improved ability to distinguish local adaptation from other models of natural selection.

## Introduction

One of the cornerstones of evolutionary genetics is understanding the processes that drive genetic differences within and among populations. A research area of particular interest is determining which observed differences in allele frequency between populations are driven by local adaptation versus genetic drift or background selection. Local adaptation is of special importance in part because population-specific selective sweeps can contribute to the initial stages of allopatric or parapatric speciation by generating genetic incompatibilities between regional variants. Furthermore, the genomic signatures of local selective sweeps can provide insights into the molecular basis and genetic architecture of adaptive phenotypic traits as they relate to regional variation.

Among the most notable and well-documented examples of local adaptation include grasses that tolerate high heavy metal concentrations in contaminated soils (Antonovics and Bradshaw 1970, Macnair 1987), mouse coat color variants adapted to different substrates by selection for camouflage (Nachman et al. 2003, Hoekstra et al 2006), the evolution of body shape and coloration of sticklebacks in response to different species of predator (Colosimo et al. 2005, Miller et al. 2015, Gygax et al. 2018), the evolution of toxin resistance in garter snakes that co-occur in regions with highly toxic newts as prey (Brodie et al. 2002), and the independent adaptation to hypoxic environments for humans and other large mammals living at high altitudes (Yi et al. 2010, Julian and Moore 2019, Witt and Huerta-Sanchez 2019). *Drosophila melanogaster* and its close relatives have also provided important model systems for genetic studies of local adaptation, including for studies focused on adaptation to high altitudes (Lack et al 2016, Sprengelmeyer and Pool 2021, Sprengelmeyer et al 2022), upper latitudes (Adrion et al. 2015; Svetec et al. 2015; Siddiq et al. 2019), and novel food sources (Yassin et al 2016).

One general approach to identifying and characterizing local adaptations is based on quantifying the differences in allele frequencies among populations at different loci, on the assumption that highly divergent loci are potential genomic signatures of local adaptation. A variety of statistical approaches have been proposed and used to leverage this information.

### Fixation Index and Population Branch Statistics

Genome-wide scans for selection identify outliers in the distribution of differences in allele frequency between a focal population and its outgroup(s) using various measures of genetic distance. Among the most widely used is Wright’s fixation index F_ST_, defined as the ratio of among population variance to the total variance across populations. The quantity was originally derived for single loci and sites, but can be generalized as a statistic for genomic regions (e.g. Reynolds et al 1983). Many studies have used high F_ST_ between two populations at specific loci in comparison to the rest of the genome or chromosome region to infer local adaptation (*e.g.* Akey et al. 2009, Amato et al. 2009, Kapun et al. 2020).

There are several drawbacks to using F_ST_ to measure population divergence, such as its non-linearity and non-additivity (making comparisons across multiple populations difficult). Importantly, when comparing very large numbers of sites in a genome-wide scan, it is inevitable that a significant fraction will be divergent among populations due to genetic drift, resulting in some fraction of loci that are falsely identified as having evolved under positive directional selection. Further, the use of F_ST_ leaves the problem of determining which population in a divergent pair has experienced local adaptation.

Particularly to address the latter problem, several statistics have been proposed that use a rescaled F_ST_-based value across population triplets (a focal population and two outgroup populations) to estimate population-specific differentiation. Among these is the Population Branch Statistic (PBS), first proposed in Yi et al (2010) as an estimate of the branch length of the focal population. The distance metric for population pairs is a log-transformed (Cavalli-Sforza 1969) function of F_ST_,

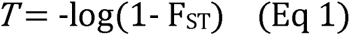

Defining a three population tree (Figure 1A) with focal population A, a relatively closely related population B, and a ‘population outgroup’ C, we can estimate the corresponding branch lengths of *a,b,c*. Because the pairwise distances *T* are approximately additive, the branch length *a* from the three pairwise *T* values is estimated as:

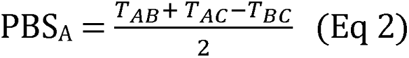

**Figure 1.**
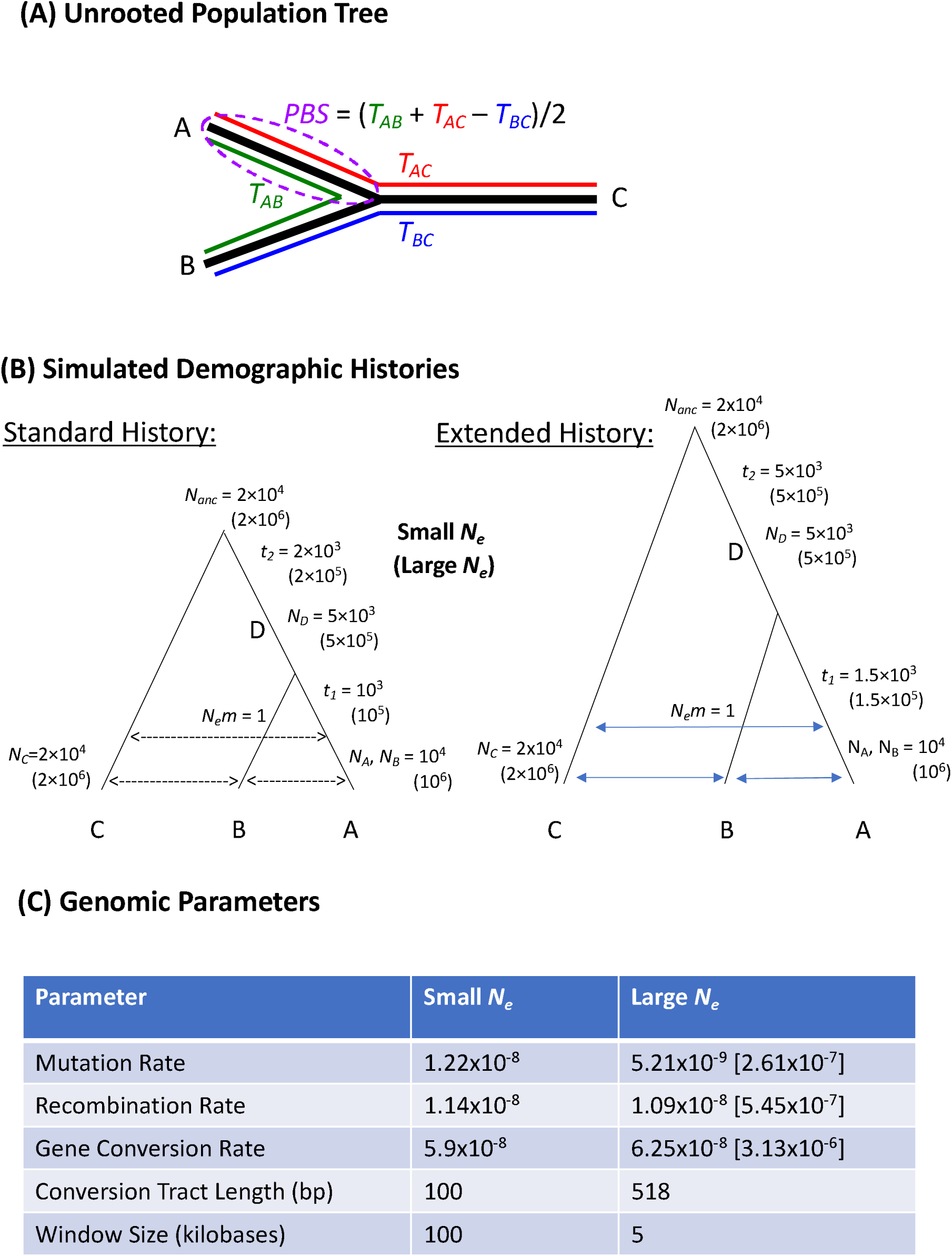
Illustration of the demographic models and genomic parameters implemented in population genetic simulations. (A) Unrooted three population tree, with A representing the the focal population. *T* values represent the genetic distances between each pair of populations, based on a log transformation of *F*_ST_ (Eq. 1). PBS, as the estimated length of the focal population branch, is then an intuitive function of these *T* values. (B) The simulated three-population genealogies. *N*_i_ represents the size of the *i*th small population, the value in parentheses is the size of the corresponding simulated large population (the lineage D is the ancestral population for A,B before their split, *N*_anc_ is the size of the population ancestral to all three sampled populations). *t*_1_ and *t*_2_ are the split times for the inner and outer divergences, respectively. The left genealogy was used for the migration-free simulations as well as the first set of migration simulations (a net population migration rate of *N*_e_*m* = 1 is represented by the dashed lines). The right genealogy has branch lengths adjusted to generate the same pairwise F_ST_ as in the first genealogy without migration under genetic drift alone. (C) The genomic parameters used in simulations for the small and large populations (values in parentheses are the 50x rescalings used in the simulations).

PBS as an estimator of focal branch length has been used in a number of recent studies to identify sites or regions of the genome evolving under selective sweeps (e.g. Jiang and Assis 2020), with the logic that loci with high PBS compared to other loci in the genome are strong candidates for selective sweeps since population divergence.

To the extent that studies aim to identify population-specific positive selection in particular, a potential disadvantage to PBS is its inability to distinguish between cases where only the focal branch is especially long from cases where all population branches are long. While sites under strong local directional selection will have high PBS, the same is true at sites undergoing similar, parallel selective sweeps in all three populations. Depending on the goals of the study, such loci may or may not be of interest, as they may often reflect instances of positive selection unrelated to adaptation to local environments, such as arms races driven by meiotic drive or reproductive competition. Furthermore, background selection (Charlesworth et al 1993, 1995) may have a relatively widespread influence on genetic diversity in at least some regions of the genome (e.g. regions with low recombination rates and/or a high density of functional sites), and frequency changes driven by background selection may also lead all populations to have unusually long branches at certain loci. Consequently, PBS may be effective at identifying instances of recent natural selection generally while being less effective at discriminating between population-specific versus species-wide selection pressures (although such predictions remain to be tested).

To address the above concerns regarding PBS, two branch statistic rescalings have been proposed with the specific purpose of identifying loci with population-specific elevations in genetic differentiation. The normalized population branch statistic, *i.e.* PBSn1, was introduced by Malaspinas et al. (2016; see supplemental section S16) and also featured in subsequent studies (Crawford et al 2017, Vicuna et al 2019). This metric rescales PBS_A_ with respect to the total tree length, e.g.

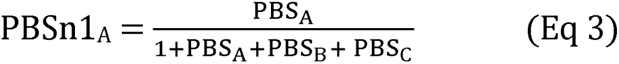

This rescaling has the potential to discriminate between cases where PBS_A_ is high in comparison to the lineages of populations B and/or C from cases where B or C also have long branches. Consequently, PBSn1 is expected to give a lower false positive rate in identifying local sweeps at loci that are evolving under parallel (*i.e.* separately occurring) selective sweeps in multiple populations. The addition of 1 in the denominator prevents division by zero and provides a baseline (although arbitrary) normalization term so that cases where all three branches are extremely short but the focal population is several times longer is not scored as significant in the same way that an especially long focal branch in particular would be (*e.g.* without the 1, a case of branch lengths *a,b,c* = 0.2,0.1,0.1 and *a,b,c* = 20,10,10 would be treated equivalently).

Unlike the above statistics, another elaboration on PBS, known as Population Branch Excess (PBE; Yassin et al. 2016), incorporates branch length information from additional genomic loci. This method posits that in the absence of local adaptation at a given locus, the relationship between the focal population’s branch length at this locus (*i.e.* PBS_A_) and the combined lengths of the two non-focal population branches (*i.e. T* _BC_) should be similar to the relationship between these two quantities observed at most other genomic loci. PBE employs this logic to obtain an expected value for PBS at this locus, based on *T* _BC_ at this locus and the median values of both PBS and *T* _BC_ observed at all other analyzed loci in the genome (or else a component of it such as the same chromosome arm). PBE then quantifies the degree to which the observed PBS value for this locus exceeds its expected value, as follows:

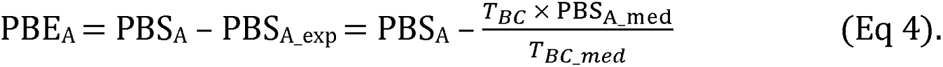

The comparison to an expected PBS value may make PBE values somewhat more comparable across studies that involve populations with very difficult demographic histories. For example, if all loci are highly differentiated due to a recent bottleneck in the focal population’s history, PBE will adjust all values based on the relatively high expected PBS (although an elevated neutral variance may still be expected). The ratio of branch lengths *T* _BC_/*T* _BC_med_ also corrects for cases where the non-focal branches are also long due to selection in those populations as well (whether sweeps or background selection). Or viewed another way, a larger-than-usual value of *T* _BC_ at this locus increases the value expected for PBS, and thus will tend to yield lower PBE values, which is desirable if the goal is a specific focus on population-specific positive selection.

We also note that multiple branch-oriented statistics have been developed to detect local adaptation from data sets that include greater than three populations (Schmidt et al. 2019; Schlebush et al. 2020; Cheng et al. 2022). However, this study’s focus is restricted to frequency-based statistics that incorporate at most three populations, in part for the sake of clarity, and in part because of the relatively larger number of empirical studies that can generate adequate population genomic data for these methods.

The heuristic considerations that prompted the proposal of PBS, PBSn1, and PBE as measures of population divergence suggest that all branch statistics may have greater power and accuracy in identifying instances of local selective sweeps against a backdrop of neutral evolution than F_ST_, and that PBSn1 and PBE may outperform PBS when some loci evolve under parallel sweeps while the evolution of other loci is driven by local adaptation. However, their relative efficacy and statistical power have not been systematically examined. Indeed, all three of the branch statistics highlighted here were introduced in the context of empirical genome scan studies which did not evaluate the performance of these statistics via simulation.

In this study, we analyze data from population genetic simulations representing a wide range of scenarios, in order to assess the above predictions regarding the performance of the above-mentioned frequency-based statistics to detect population-specific positive selection. We include simulations with large and small population sizes (motivated by *Drosophila* and humans, respectively), and we investigate models with and without gene flow after population divergence. We combine simulated replicates into simulated genomes, and assess which scenarios are placed in the top (upper 1%) outlier quantile for each statistic, emulating an empirical outlier genome scan. This approach allows us to differentiation local adaptation not only from neutral evolution but also from background selection (modeled as reduced population size) and parallel selective sweeps. Results of these analyses will allow researchers to make more informed choices regarding which statistics to use in future population genomic scans for local adaptation.

## Results

We performed population genetic simulations to test the ability of four statistics (F_ST_, PBS, PBSn1, and PBE) to detect population-specific positive selection, examining three population models in small and large *N*_e_ cases motivated by human and *Drosophila* data, respectively. The demographic and genomic parameters used in these models are summarized in Figures 1B-C. We aggregated simulations from distinct neutral/selection scenarios into “model genomes” (Figure 2A) to test how reliably each statistic placed the 1% of true local sweep loci in its the upper 1% tail, with or without an additional 1% of loci subject to parallel sweeps in all three populations, and with the remaining loci subject to either neutral evolution or background selection (BGS; modeled as variably reduced *N*_e_).

**Figure 2.**
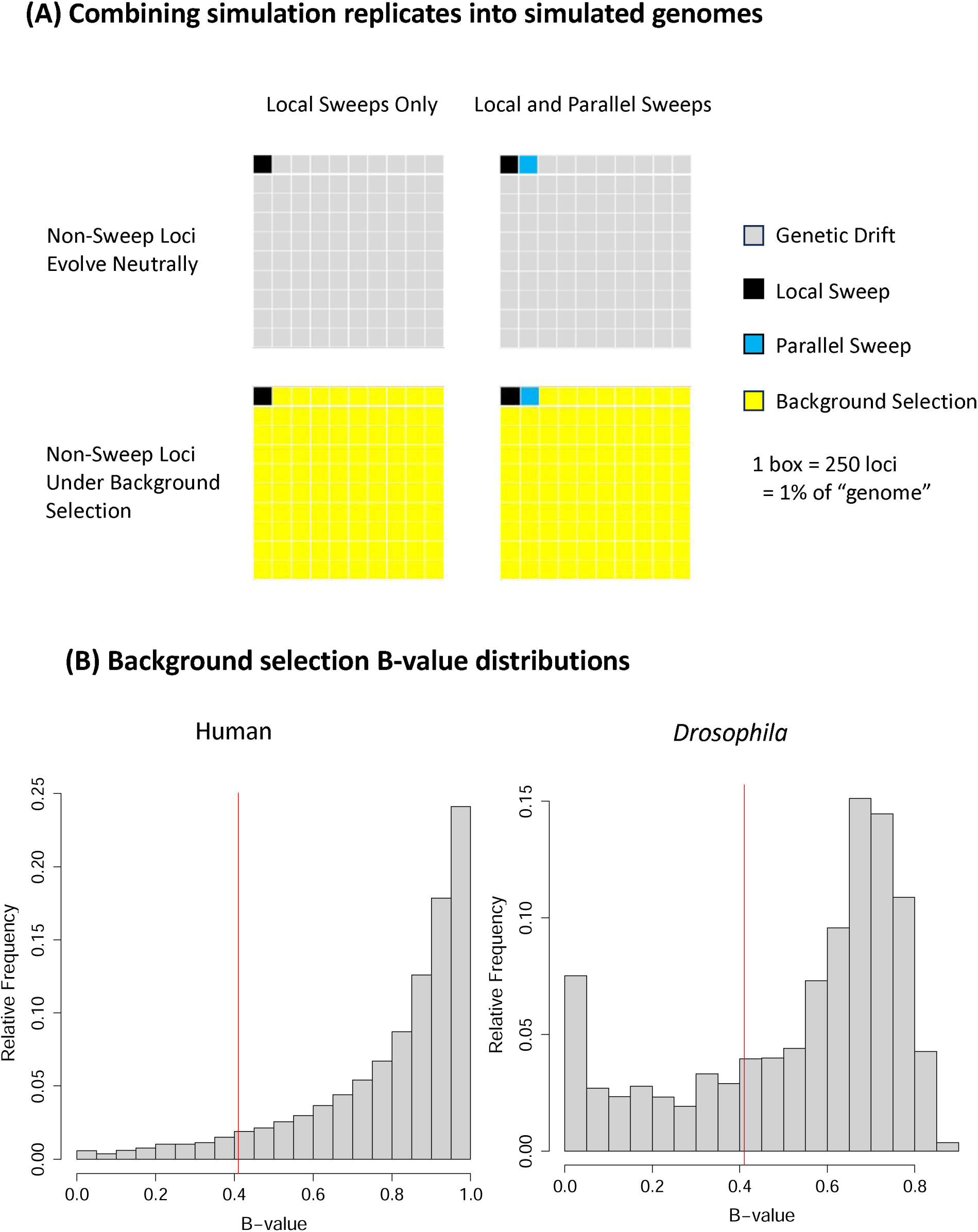
Simulated genome models used in this study. (A) Schematics representing the model genomes each containing 25,000 simulated loci (representing genomic windows in an empirical scan). The upper left genome has 99% of loci randomly sampled from the 10^6^ genetic drift simulations and 1% from the 10^4^ local selective sweep simulations. The upper right genome has 98% neutral, 1% local selective sweep simulations, and 1% parallel sweep simulations. The lower left genome has 99% BGS loci and 1% local sweeps. The lower right genome has 98% BGS loci, 1% local sweeps, and 1% parallel sweep loci. (B) The autosomal B-value distributions for human (left) and *D. melanogaster* (right) genomes are shown, as used here for the simulations of small and large population simulations, respectively. The vertical red line rep resents the truncation at *B* = 0.41for each population size scenario, in order to approximate genome scans in which low recombination regions are excluded due to the difficulty in localizing targets.

We simulated local complete hard sweeps with selection coefficients *s* = 0.025 and 0.001 in the small and large populations, respectively. When these local sweeps occur amongst neutral loci, the loci under selection can be identified with a high level of accuracy by all statistics, with only slight advantages of the three population statistics over pairwise F_ST_. As we deviated from this simple scenario, the statistics began to diverge from one another in both their precision based on the model genomes defined above (Figure 3) and their more traditional statistical power based on comparing individual local sweep replicates to neutral or BGS distributions (Table S1).

**Figure 3.**
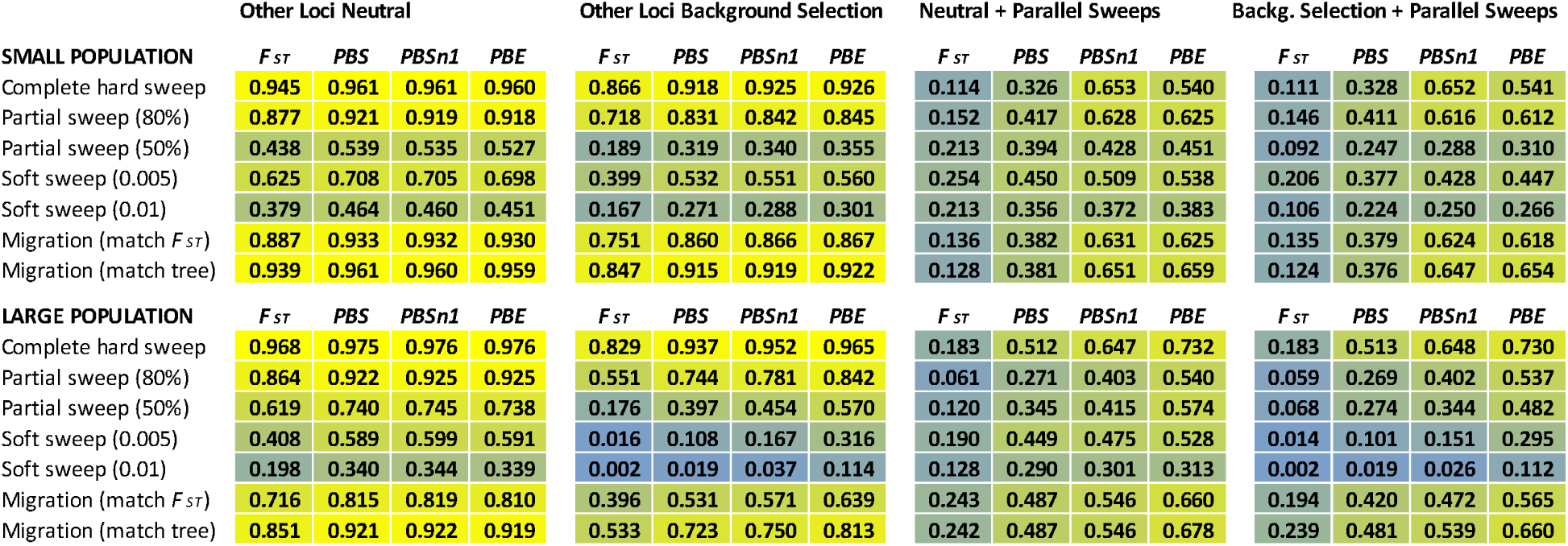
Population branch statistics show greater precision to detect local selective sweeps than F_ST_ when other loci experience global positive or negative selection. Heat maps show the precision of F_ST_ and the population statistics PBS, PBSn1, PBE with respect to local sweeps, i.e. the fraction of local sweep loci among those contributing to the upper 1% quantile of each statistic. The table includes all demographic scenarios (large and small population, with and without migration), genomic backgrounds (genetic drift vs. BGS), and selection regimes (hard complete, partial, soft complete) considered in the study. Migration scenarios involved hard sweeps that might have fixed if not for gene flow.

Precision was reduced in the cases of partial and soft local sweeps in otherwise neutral genomes, with PBS, PBSn1, and PBE often performing similarly, and F_ST_ consistently showing the lowest precision (Figure 3). In contrast, in model genomes where most loci evolve under BGS rather than genetic drift, the relative performances of the branch statistics became more distinct: here, the precision of PBSn1 and PBE was invariably larger than for PBS in both the large and small populations (Figure 3). PBE also outperformed PBSn1 in each of these cases, although the difference was more notable in the large populations. In the scenario that yielded the most disparate outcomes, that of a soft sweep from 0.5% initial frequency in a large population with BGS, precision was 1.6% for F_ST_, 10.8% for PBS, 16.7% for PBSn1, and 31.6% for PBE. In a few of the most challenging cases, the precision of some or all statistics fell considerably. In the large population case, strong effects of the examined BGS model even led some precision estimates to fall below the null expectation of 0.01, e.g. for a soft sweep with *p*_0_ = 0.01 (perhaps partly because the sweep loci themselves were not subjected to BGS). Under these scenarios, BGS tends to generate longer branch lengths than genetic drift (especially in the large populations, which have a higher fraction of loci with *B* < 0.5; Figure 2B). Here, PBE’s rescaling of branch lengths with respect to all loci in the “genome” may improve its performance in comparison to PBS and even with respect to PBSn1, which rescales the focal branch length with respect to total tree length at the focal locus but not with respect to an overall genome-wide baseline.

In simulations that included both parallel and local sweeps at 1% of the loci, in otherwise neutrally evolving genomes, the overall frequency of false positives increased for all of the statistics when compared to otherwise equivalent scenarios in which all sweeps were local (Figure 3). Here, the advantages of PBE and PBSn1 over PBS became more pronounced, with PBE again showing a larger advantage for the large population scenarios. Whereas, the performance of F_ST_ became especially poor with parallel sweeps. In these scenarios, we expect that parallel sweeps may generate higher F_ST_ values than local sweeps, so parallel sweeps will disproportionately contribute to the upper quantiles of F_ST_ and potentially PBE (which simply looks for a long estimated focal population branch length). The enhanced differentiation of parallel sweeps (and thus their contribution to false positives) can be greater for hard complete sweeps than for partial or soft sweeps (Figure 4), which may contribute to the otherwise surprising observation that in some cases, F_ST_ and to lesser extent PBS had lower precision and power with hard complete sweeps than with certain soft or partial sweeps. This phenomenon is less apparent in the large populations, where the modeled selection coefficients are weaker and the rate of recombination relative to mutation is somewhat higher (see Materials and Methods).

**Figure 4.**
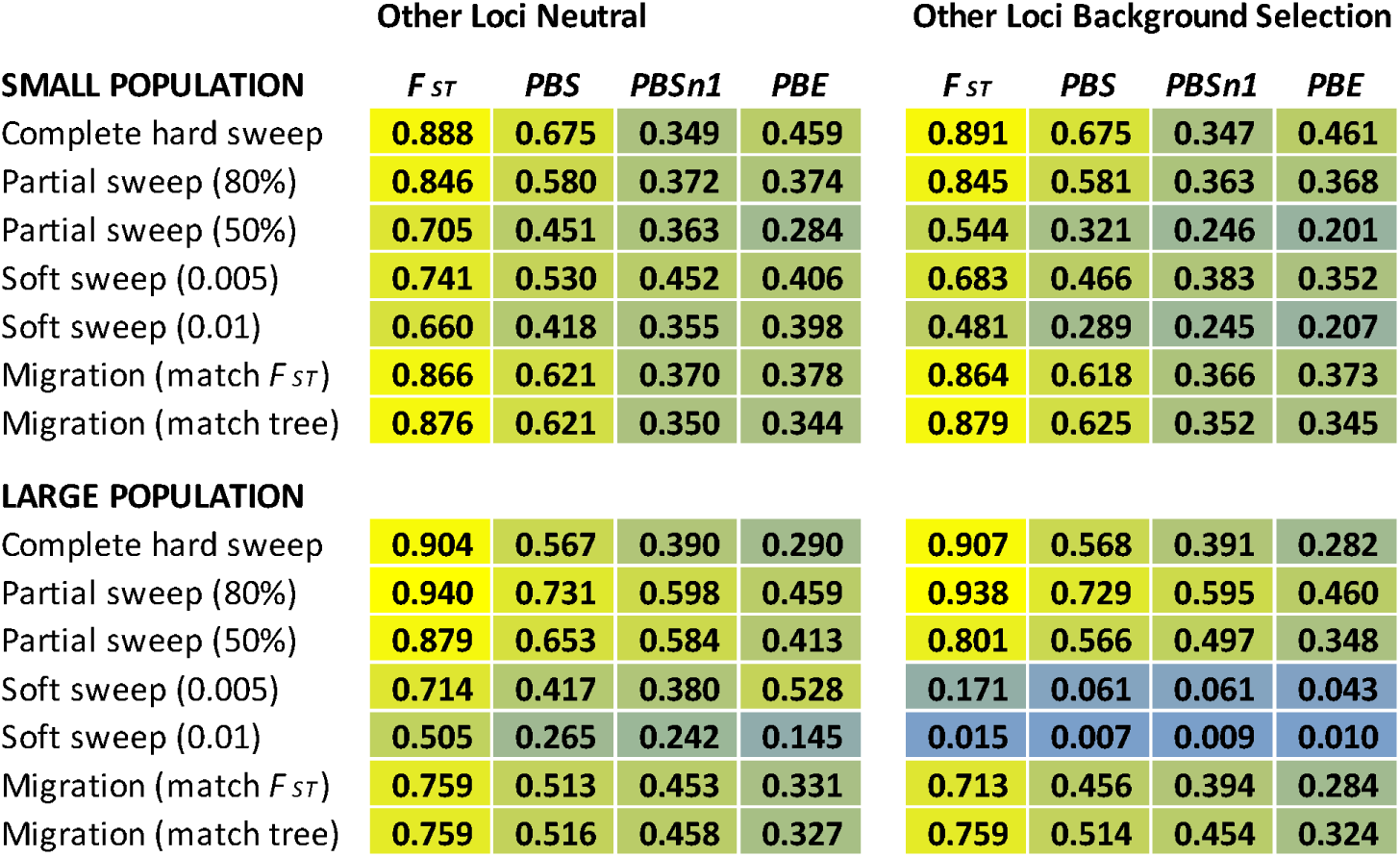
PBE and PBSn1 are less likely than other statistics to register parallel sweep loci among their top outliers. For model genome scenarios in which 1% of loci are subject to parallel sweeps in all populations, the heat maps show the fraction of parallel sweep loci that contribute to the upper 1% quantile of F_ST_ and branch statistic distributions (i.e. the fraction of parallel sweep loci that are false positives) under different demographic and selection parameters.

With hard complete local and parallel sweeps (as well as partial sweeps to 80% terminal frequency), most false positives were contributed by parallel sweep loci (Fig. 4, Table S2), while for soft sweeps at high initial frequency and for partial sweeps to 50%, there was a greater contribution from the neutral and BGS loci (especially for the case of BGS in the large populations – e.g. 46-94% of parallel sweep loci contributed to upper quantiles under 80% partial sweeps whereas only about 1% of parallel sweep loci were in the upper quantile in soft sweeps with *p*_0_ = 0.01). The proportional contribution of parallel sweeps vs. neutral/BGS loci to false positives impacts the relative ability of the population statistics to detect local sweeps. For example, in a small population with both local and parallel sweeps, the precisions for some of the statistics, especially F_ST_ and PBS, are higher for soft sweeps (with *p*_0_ = 0.005) and partial sweeps (*p*_t_ = 0.80) than for complete hard sweeps. This counter-intuitive pattern is not seen in the large population simulations, which have lower false positive rates from the parallel sweeps than in the small populations. In contrast, with very soft sweeps (*p*_0_ = 0.01) or partial sweeps to only 50% final frequency, there is also a significant relative contribution of neutral or BGS sites to the false positives.

When populations are subject to both parallel sweeps and background selection, similar trends hold, with slightly to moderately reduced precision for each statistic to detect local sweeps compared to cases with parallel sweeps without BGS. In all cases, F_ST_ showed the poorest performance, while PBS lagged behind the other two branch statistics. For small populations, PBSn1 and PBE show similar advantages, while for large populations, PBE displayed consistently greater precision, generally showing similar or greater advantages over PBSn1 and the other statistics than observed with either parallel sweeps or BGS alone. For example, in the case of soft sweeps with *p*_0_ = 0.005, PBE at 0.295 gave a precision nearly two-fold and three-fold higher than PBSn1 and PBS, and twenty-fold higher than F_ST_.

While the above-mentioned results all pertain to simulations with genetic drift but no migration, we also explored cases in which all pairs of populations have exchanged one migrant per generation (*i.e. N*_e_*m* = 1) since their divergence. Adding this level of migration to the previously studied population history had a relatively weak effect on the precision and power of the statistics to detect hard local sweeps under scenarios lacking any other form of selection (Figure 3; Table S1; “match tree”). Here, while migration may have reduced the frequency differences at locally adaptive loci, it also resulted in populations with lower neutral genetic differentiation than in migration-free base scenario. The effect of migration on precision and power is more perceptible under the scenarios in which we extended the divergence times of our populations to emulate the neutral F_ST_ values observed among populations without migration (Figure 1B), since here allele frequency differences should be reduced at local sweep but not neutral loci compared to the base history (Figure 3; Table S1; “match F_ST_”). While the effects of migration on its own were not dramatic for either population size or population tree length, its effects synergized with BGS to reduce precision, particularly for the large population size with its more powerful modeled BGS effects (Figure 3). With parallel sweeps, the performance of F_ST_ and in some cases PBS actually improved with the introduction of migration. To some extent, migration may be blunting the propensity of parallel sweeps to generate false positives (Figure 4; Table S2), analogous to the outcomes of some partial or soft sweep models discussed above. In general with migration, as observed otherwise, PBE and PBSn1 showed the greatest precision when other types of selection were present, and PBE in particular showed greater advantages in the large populations.

## Discussion

Population branch statistics were introduced as alternatives to F_ST_ in genome-wide scans for local selective sweeps on heuristic grounds, without explicit assessment of their performance. The present study provides simulation-based evidence favoring their use over F_ST_ as well as supporting the use of rescaled branch statistics PBSn1 and PBE rather than PBS for a wide range of evolutionary models. Generally, only in the most trivial scenario of a small number of loci experiencing local selective sweeps while all others evolve neutrally do F_ST_-based scans or PBS have precision and power comparable to the rescaled population branch statistics PBSn1 or PBE. Otherwise, the rescaled branch statistics were consistently more robust in contributing to the upper quantiles when potentially confounding selective processes such as BGS or parallel sweeps were introduced.

This study also provides qualified support for the use of PBE over PBSn1 under many selection regimes, including the potentially more biologically realistic models of partial and soft sweeps, and in genomes where background selection plays an important role in shaping genetic differentiation. On the other hand, there are certain scenarios, such as those where there are strong parallel as well as local sweeps, where PBSn1 performs as well as or slightly better than PBE, though it is difficult to generalize from these cases because the differences are typically modest and contingent on multiple parameters. Furthermore, complete, hard selective sweeps are probably quite rare, at least in human populations (Hernandez et al 2011), which may limit the importance of the only case in which PBSn1 had a non-trivial advantage over PBE (*i.e.* both complete hard local and parallel sweeps in small populations). One factor that may modulate the relative precision PBSn1 compared to other statistics is the arbitrary scaling factor of 1.0 in the denominator (Eq 2), which may serve to dampen noise from excessively small denominators. Conceivably, the precision of PBSn1 relative to PBE might vary if other values of this scaling factor were used. On the other hand, it is worth emphasizing that PBSn1 (like PBS and F_ST_) can be calculated using only genetic variation at a specific locus, whereas each value of PBE necessarily draws on information from other comparable loci.

Given the lack of prior results, our study represents a useful starting point in evaluating the relative performance of the statistics examined. Future studies could use demographically expanded simulations to incorporate methods that accommodate more than three populations (e.g. Cheng et al. 2022). It would also be instructive to investigate a wider array of demographic models, such as strong population bottlenecks, additional migration rates, or admixture models, to further assess the context-dependency of the differences in precision we observed among statistics.

Lastly, while our use of reduced population size as a proxy for the effects of BGS on allele frequency differentiation is reasonable in the context of this study, it will be worthwhile to ultimately supplement our analyses with explicit simulations of negative selection at linked sites (Ewing & Jensen 2016). Such simulations may be especially computationally demanding for *Drosophila* -like cases, in light of the chromosomal scale that may be relevant (*e.g.* Comeron *et al*. 2014) and the unclear effects of population size rescaling (as often conducted for simulations with *Drosophila* -like *N*_e_) on the outcomes of BGS simulations. Relatedly, while our simulation results suggested a strong potential of background selection to generate outliers in F_ST_-based statistics in the *Drosophila* -like simulations, it is worth noting that Booker et al. (2020) found that the empirical relationship between crossing-over rate and F_ST_ between *D. melanogaster* populations was roughly similar to that expected under a simulated model without background selection, particularly above the female crossing-over rate threshold of 1 cM/Mb that our B-value cutoff approximates.

Even in an era featuring increasingly complex population genetic methods for detecting local adaptation, we suggest that there is value in simpler statistics that have relatively straightforward interpretation with regard to genetic variation. In this context, we suggest that rescaled population branch statistics such as PBE and PBSn1 are relatively powerful and efficient tools for identifying the genetic signatures of local adaptation in genome-wide scans. The results of this study argue in favor for their expanded use in the toolkit of those seeking evidence of local adaptation in natural populations, particularly in cases where other forms of positive or negative selection may impact genetic differentiation. Furthermore, these statistics and their underlying logic may continue to provide useful building blocks for more advanced methodologies in the future.

## Methods

We simulated evolution in a three-population model where the evolutionary dynamics of genotype frequencies are driven by (at different loci in various combinations) neutral genetic drift, background selection against deleterious mutations, local selective sweeps, and parallel selective sweeps in all three populations. Unless otherwise indicated, simulations were run using Python 3.8.8 scripts and all data analyses were performed using R 4.1.1.

### Population Demographic History

We chose two base demographic models to approximate the population histories of humans and *Drosophila melanogaster* as representatives of large and small population species with respective effective sizes of *N*_e_ ∼ 10^4^ and 10^6^. We set the outer and inner population split times at 2 × 10^3^ and 10^3^ generations before present for the small population case and 2 × 10^5^ and 10^5^ generations for the large population case (as shown in Figure 1B). These time intervals roughly approximate the split times between Africa vs. Eurasia and Eurasia vs. the Americas for humans and between sub-Saharan Africa vs. the Mediterranean and the Mediterranean vs. north-central Europe for the flies.

We applied a reduced population size for the internal branch representing the common ancestor of populations A and B (labeled in Figure 1B as population D, which is half the size of its descendant populations), in order to roughly recapitulate the genetic drift involved in both of the motivating species’ founder event expansions from Africa into Eurasia (note that this bottleneck was much stronger in human evolutionary history). Note that we are not attempting to model the precise demographic details of either species’ history, rather, we use these approximate and simplified topologies to capture the qualitative differences between large and small population models.

To make large population forward simulations computationally feasible in light of the very high number of generations needed, we modeled evolution using a rescaling approximation where a 50-fold smaller *N*_e_ population was simulated in combination with 50-fold higher rescaling of selection coefficients, mutations rates, and other populations parameters, an approach similar to the one used by Lange et al. (2018). The demographic and genetic parameters are listed in Figure 1C for both the small and large populations (based on ancestral *N*_e_ = 2 x 10^4^ and 2 × 10^6^, respectively, simulated with a corresponding 50x rescaling in the large population). We used this rescaling in both the forward time (selection) and backward time (coalescent drift) simulations for consistency when simulating evolution in the large populations.

### Parameterizing the Model Genomes

We generated genomic windows so that in our simulations a “locus” is defined not as a single site, but as a chromosome region. We simulated 100 kb and 5 kb regions for the small and large populations, respectively, following previous simulation studies (*e.g.* Lange and Pool 2016; da Silva Ribeiro *et al*. 2022), and equal or close to window sizes from empirical genome scan studies (*e.g.* Voight *et al*. 2006; Granka *et al*. 2012; Lack *et al*. 2015; Pool *et al*. 2017). The 20x difference in window length also represents a compromise between the approximately 10x higher population mutation rate and the roughly 40x higher population recombination rate estimated for fruit fly versus human populations. Per-site mutation, recombination, and gene-conversion rates were based on the mean autosomal estimates in *D. melanogaster* from Comeron et al (2012) and for humans from Schiffels and Durbin (2014), Williams et al (2015), and Jonsson et al (2017). Figure 1C summarizes the genomic parameters used in the simulations.

### Simulating Genetic Drift

In purely neutral simulations, genetic drift was modeled as a backward time coalescent process using the msprime package 1.2.0 (Baumdicker et al. 2022) for the demographic models and parameters summarized in Figure 1B, 1C. The msprime commands were embedded in python script “wrappers” that calculated allele frequencies and the population statistics.

At the termination of each simulation run, we calculated F_ST_ among subpopulations based on the Reynolds et al. (1983) estimate of the fixation index as well as PBS and PBSn1. The branch length *T* _BC_ was also output for each simulated replicate, since this is needed alongside PBS values for the subsequent calculation of PBE, which is calculated with respect to an aggregate of loci in a genome rather (Eq. 4). Additionally, sequences of segregating sites generated by the simulations were saved as output.

### Background Selection

The effects of background selection (Charlesworth et al. 1993, 1995) were simulated without explicitly modeling deleterious mutations or negative selection as such. Rather, we modeled BGS implicitly by simulating the effects of negative selection on the genetic diversity of linked neutral variation. Charlesworth (2012) demonstrated that the reduction in linked neutral genetic variation due to BGS is equivalent to that under a reduction of effective population size by a factor *B* which depends on the recombination rate in a genomic window (i.e. regions of low recombination correspond to B-values much less than one). As examples, rescaling *N*_e_ (and therefore expected coalescent times) with B-values to model the effects of BGS on neutral genetic variation was applied by Jiang and Assis (2002) to estimates of F_ST_ among populations evolving with BGS, and in the calculations of Huber et al. (2015) of allele frequency spectra under BGS. There is evidence that modeling BGS as drift with reduced *Ne* is inadequate for demographic inference (Ewing and Jensen 2016), and it would seem similarly inappropriate for studies making quantitative inferences about the background selection process itself. In contrast, because the B-value approximation does capture the reduction in nucleotide diversity due to BGS, it should recapitulate the resulting increases in allele frequency differentiation between populations to a sufficient degree for a study not focused on parameter estimation.

Specifically, we model the effects of BGS on neutral genetic variation by using the distribution of B-values estimated for chromosome regions in *D. melanogaster* in Comeron (2014) and for the human genome in McVicker et al. (2009). In each simulation, a value of *B* is randomly drawn from the frequency distribution (from the *Drosophila* autosome distribution for large population simulations, the human autosome distribution for small populations) and used as a multiplier to rescale the effective population size. The evolution of genetic diversity in a region of genome with this value of *B* is simulated as genetic drift using msprime with *N*_e_* = *BN* _e_ in place of *N*_e_.

Certain chromosome regions, particularly those near centromeres in *Drosophila*, have very low *B*, i.e. < 0.01 (Figure 2B), resulting in very small rescaled effective population size and greatly inflated F_ST_ for simulated loci in those intervals. We corrected this potential artifact by removing B-values corresponding to regions where the sex-averaged crossing-over rate *r* < 0.5 cM/Mb, analogous to the removal of low recombination regions such as those around autosomal centromeres in many genome-wide scans of *D. melanogaster* (*e.g.* Pool et al. 2017). Low recombination regions are often excluded from genome scans because of (1) the greater variance in diversity patterns expected under neutrality (Booker et al. 2020), (2) stronger effects of background selection, and (3) the difficulty in localizing outliers due to positive selection in low recombination regions.

To find an appropriate cut-off, we performed a LOESS regression of *D. melanogaster* B-values against recombination rate and identify the value of *B* predicted at *r* = 0.5 by the regression model. This corresponds to *B* = 0.41, so the B-value distribution was truncated at *B* > 0.41. For consistency, this was done for both the human and fruit fly distributions, even though human chromosomes had only 7% of regions where *B* < 0.41 versus 30% of fruit fly autosomes. Simulations of BGS as genomically variable drift were run by sampling *B* from these truncated distributions of B-values for each simulated genomic window (*i.e.* simulation replicate).

### Selective Sweeps

All simulations of positive selection were performed using the SLiM v. 3.5 software package (Messer 2013, Haller et al. 2019). Unlike the coalescent-based models in msprime, SLiM simulations run in forward time. The SLiM output consists of a tree structure (the sequences and their genealogical history) of the site(s) under selection. To generate the corresponding history of neutral sites that evolve in association with the sites evolving under directional selection, we used msprime to simulate their coalescent history and link them to the tree structure generated in SLiM, a process known as recapitation of the genealogy, as described in Kelleher et al. (2018) and Haller et al. (2019).

Most of the SLiM simulations were initiated by introducing a beneficial mutation at a randomly selected single site in the focal population in the first generation following its split from its outgroup. The simulation runs were conditioned on either the fixation of the beneficial allele or it reaching a specified terminal frequency *p*_t_ < 1.0. If the beneficial mutation was lost from the population during the simulation, the simulation was terminated and restarted. Thus, only the output tree structures of those replicates that reached the desired terminal frequencies were retained.

Variations on this basic model of selective sweeps included conditioning on intermediate *p*_t_ < 1.0 (partial sweeps), initial frequencies orders of magnitude greater than 1/2*N* (soft sweeps), and simultaneous selection at the same locus in both the focal population and its outgroups (parallel sweeps). All of the selective sweep scenarios described below were run to generate 10,000 replicates for analysis.

#### A. *Complete local hard sweeps*

These simulations used selection coefficients *s* = 0.025 and 0.001 in the small and large populations, respectively (with the smaller *s* in the larger *N*_e_ populations reflecting the expectation that weaker beneficial mutations can successfully be favored by selection in larger populations). With the population size rescaling by 50x of the large populations, the rescaled selection coefficient used in those simulations was *s*’ = 0.05.

Because of the high probability that a single beneficial mutation will be lost or fail to be fixed in a simulation with small *N*_e_ and *s* << 1, we used an initial frequency *p*_0_ = 0.001 rather than *p*_0_ = 1/2*N*_e_ (set at the time of the focal population’s split from its outgroup) in the small *N*_e_ scenarios, to reduce the number of times that simulations had to be restarted to achieve fixation. However, in the large *N*_e_ scenarios, we used *p*_0_ = 1/(2*N*_e_) to avoid soft sweep effects in the rescaled model populations. Previous studies, e.g. da Silva Ribeiro et al. (2022) have shown that these initial frequencies are sufficiently low to generate hard sweep outcomes (*i.e.* only one haplotype carrying the beneficial haplotype tends to contribute to the sweep). Additionally, because many simulations didn’t reach fixation by the final generation, rather than restarting all simulations where the *p*_t_ < 1.0, we implemented a strategy in which simulations with *p*_t_ > 0.95 were saved and only those haplotypes with the beneficial mutation were sampled for F_ST_ estimation and other analyses. These near-fixations provide a close approximation to a complete hard sweep, and the approach was also used when modeling parallel and soft sweeps.

#### B. *Complete local soft sweeps*

Soft sweeps can occur when positive selection acts on alleles that had already achieved a frequency significantly higher than 1/(2*N*_e_) in the population through genetic drift, and multiple initial haplotypes contribute to the sweep, contributing to greater haplotype diversity after fixation than hard sweeps (Hermisson and Pennings 2005). We simulated soft sweeps with initial beneficial allele frequencies of *p*_0_ = 0.005 and 0.01 in both the large and the small populations, using the same selection coefficients and initiation times at the A,B split as in the complete sweep simulations.

#### C. *Partial hard sweeps*

In practice, many beneficial alleles may never become fixed, or they may be sampled at time points prior to their fixation in a population. To simulate (hard) partial sweeps, we applied positive selection to an allele in the focal population using the same initial frequencies and selection coefficients as described for complete local sweeps. To target sweeps reaching terminal frequency *p*_t_ = 0.5, we introduced the beneficial mutations at *t* = 1700 and 3800 generations, i.e. 300 and 200 generations prior to termination in the small and large populations, respectively; for *p*_t_ = 0.8, the beneficial mutations were introduced at generation times 1500 and 3700. These initiation times were chosen because they generated terminal frequencies close to the desired values. Simulations were repeated until a terminal allele frequency between 0.45 and 0.55 was observed in the last generation to simulate partial sweeps to 50% (or between 0.75 and 0.85 to approximate *p*_t_ = 0.8). Unlike for the complete sweeps, haplotypes were sampled randomly from the entire population at the end of each saved simulation, as opposed to only sampling haplotypes carrying the beneficial allele.

#### D. *Parallel sweeps*

We simulated parallel selective sweeps by introducing a distinct beneficial mutation in all three populations at the same position. For hard complete parallel sweeps, we used the same initial frequencies and selection coefficients as were used for hard complete and partial local sweeps. Beneficial mutations were introduced at *t* = 1000 and *t* = 2000 at the same locus in all three populations (corresponding to the time of the A,B population split in the small and large populations, respectively) in order to allow maximum time to reach fixation. As with hard complete local sweeps, the simulations were re-run until the beneficial mutations reached *p*_t_ > 0.95, and if not fixed, only haplotypes carrying the beneficial mutation were sampled for analysis.

We also simulated soft parallel sweeps and partial parallel sweeps so that the resulting allele frequencies and F_ST_-based statistics could be compared to those generated by the same positive selection models in local sweeps. The parallel soft and parallel partial sweeps used the same initial frequencies, selection coefficients, initiation times for the introduction of beneficial mutations, and targeted terminal frequencies as in the corresponding local sweep simulations, but applied these conditions to all three populations rather than the focal population alone.

### Simulating Migration

A subset of simulations included migration among all populations, in order to determine the extent to which results on the relative efficacy of branch statistics were robust in the presence of gene flow. We considered two demographic models with migration. One retains the split times of the migration-free models (left tree in Figure 1B), with a symmetric pairwise migration rate introduced so that *N*_e_*m* = 1 per generation between all pairs of populations. The migration rates were set so that each population received on average a single migrant per generation independent of population size (such that *m* was smaller for small population receiving migrants from a large population than the reverse) to avoid source-sink effects. In the rescaling of the large populations, we set *N*_e_*’m’ = 1* (where *N*_e_*’ = N*_e_*/50*, *m’* = 50*m*) so that there would be comparable numbers of migrants exchanged among pairs of populations over the same number of coalescent time units.

Because migration is expected to reduce pairwise differentiation among populations, the baseline expected F_ST_ at neutrally evolving sites will be lower in the presence of migration than in its absence, so a high or low branch statistic in the absence of migration may not be comparable to values observed in populations evolving with migration. Therefore, we also implemented a longer branch demographic history for a three population model with migration that generates very similar pairwise F_ST_ among all pairs as the migration-free model. For the same population sizes, we introduced split times at 3×10^3^ and 5×10^3^ generations before the present for the small population models and 3×10^6^, 5×10^6^ for the large population models (right tree in Figure 1B).

Simulations of genetic drift and selection with migration were executed in msprime and SLiM with the same selection coefficients in the small and large populations as for the models of selective sweeps as in the models without migration.

Because of migration, the selective sweeps are not expected to be complete, so we conditioned the local sweeps on attaining a terminal frequency *p*_t_ > 0.75, and imposed the same constraint on all three populations when simulating parallel sweeps in both the large and small *N*_e_ scenarios. This threshold value was selected to eliminate cases where the beneficial mutation was nearly lost while allowing for intermediate frequencies due to migration (*e.g.* when simulating 1000 replicates of local hard sweeps, we found that 98% of the replicates attained frequencies *p*_t_ > 0.80). As with the partial sweep simulations, all haplotypes were included in the final sample rather than only those carrying the beneficial mutation because we are not simulating fixation. Beneficial mutations were introduced at the time of the split between populations A,B, as was done for the hard complete sweep models without migration.

In simulations of migration with local sweeps, the beneficial mutation in the focal population was neutral in the two non-focal populations. Similarly, when modeling parallel selection with migration, the beneficial mutation for each population was neutral with respect to the wildtype background in the other two populations (and thus relatively deleterious as a migrant, compared to another beneficial mutation within its home population).

### Model Genomes and Statistical Analysis

The msprime simulations, including the recapitated SLiM trees, generate a distribution of multilocus genotypes from which we calculated allele frequencies in the three populations, and F_ST_ at each locus for every population pair. We used the F_ST_ estimator in Reynolds et al (1983) and Weir and Cockerham (1984) because it incorporates the contribution of sampling variances and generalizes to a multilocus estimator for F_ST_ across multiple sites, allowing F_ST_ to be calculated for genomic “windows”. The branch statistics PBS, and PBSn1 for focal population A were calculated from the genomic window F_ST_ using Equations 1 and 2.

Here, each model genome consisted of a collection of simulated replicates that include both a subset of local sweep replicates as well as replicates from one or more other evolutionary models. We created four types of model genomes representing scenarios where the majority of sites evolved either neutrally or under BGS, while a small fraction experienced selective sweeps (Figure 2A). The genomes consisted of 25,000 loci (genomic windows from independent simulation replicates) for both the small and large populations, roughly aligning with the full genome size for human and *Drosophila*, respectively. In one scenario, 99% of the loci were sampled from the million replicates of genetic drift simulations, and the remaining 1% from the 10,000 local sweep runs. A similar model had 99% of loci generated under a BGS model rather than drift (we do not simulate cases that are mixtures of BGS and drift, because our BGS models include a subset of sites with *B* close to 1 that essentially evolve as under neutral drift). The other sets of model genomes combined parallel and local sweeps, with 98% of loci sampled from the genetic drift (or BGS) replicates, 1% from local sweeps, and 1% from parallel sweeps. Focal population PBE at each locus was computed from the locus-specific PBS and *T* _BC_ and from the median PBS and *T* _BC_ values across all loci (Eq. 3) within the model genome.

After the completion of each simulation, we sampled 25 diploid individuals from each population and calculated pairwise F_ST_ and the population branch statistics. We quantified the relative performance of F_ST_, PBS, PBSn1, and PBE by calculating their probability of identifying local sweep loci. In this context, true positives were defined as local sweep loci in the upper 1% quantile of a statistic’s distribution, while false positives were any non-local sweep loci in this quantile (*i.e.* loci evolving under genetic drift, BGS, or parallel sweeps). The ratio of true positives to all positives defined the statistical *precision* of each test. For the genomes with both local and parallel sweeps, we also calculated the fraction of parallel sweep loci that contributed false negatives, and the fraction of false negatives that were from parallel sweeps rather than drift or BGS.

Above, we describe the estimation of false positive rates in a simulated genome context. As a complement to this approach, we also computed the statistical power of F_ST_ and the branch statistics in a simpler and more conventional sense. Here, we separately compared each simulated local sweep replicate against the full set of replicates simulated under a given null model (either genetic drift or BGS). The statistical power was calculated as the fraction of selective sweep replicates that were within a given statistic’s upper 1% quantile for the neutral or BGS replicates.

## Availability of Data and Materials

All of the msprime and SLiM scripts used to generate simulated data, along with scripts used to analyze that data, are available at https://github.com/mshpak76/PBE.

## Acknowledgments

The authors thank the members of the Pool lab, Josep Comeron, and Aaron Ragsdale for helpful discussion. We also thank Ben Haller, Peter Ralph, Jerome Kelleher and other developers of SLiM and msprime in helping to resolve initial issues with the simulations. This work was supported by NIH award R35 GM136306 and by NSF award DEB 1754745 to JEP.

## Author Contributions

MS and JEP created the study design and wrote the manuscript, MS and KNL ran the simulations and analyzed the output. MS and JEP wrote the paper.

